# Increased flagellation enhances bacterial migration against and across flow

**DOI:** 10.64898/2026.10.16.759531

**Authors:** Farhad Javi, Seiga Yanagisawa, Navish Wadhwa

## Abstract

Rheotaxis enables bacteria to swim against fluid flow, influencing their dispersal and colonization of surfaces. Although flagella generate the propulsion required for swimming, whether having more flagella improves rheotaxis remains unclear. Here, we varied flagellation using an *Escherichia coli* strain with tunable flagellar expression and tracked near-surface swimming under controlled microfluidic flow. Greater flagellation increased swimming speed without flow, reduced downstream transport under flow, and increased the fraction of tracks moving upstream. More flagellated cells also swam upstream for longer before switching direction. Greater flagellation also enhanced directed migration across the flow, suggesting that flagellar number alters the balance between propulsion, upstream alignment, and chiral turning near surfaces. A near-surface swimming model, in which flagellar number affects the swimming speed and the tilt of the cell toward the surface reproduced the trends in downstream transport, upstream swimming, and lateral migration. These results identify flagellar number as a determinant of how bacteria resist downstream transport and migrate across near-surface flows.

## Introduction

Swimming bacteria rarely move through still fluid. In the urinary tract, the bloodstream, catheters, soil pores, and water pipes, they swim within flows that reorient them and carry them down-stream [1–6]. Near surfaces, bacteria can resist downstream transport by reorienting and swimming against the current, so that outward flow does not always prevent contamination of ducts and medical devices [4, 7, 8]. Such upstream invasion is led by cells that sustain long runs against the flow [8], so the outcome depends not only on the direction in which cells swim but also on how long they maintain it. The reorientation and upstream swimming of cells in the presence of gradients in fluid velocity is called rheotaxis [9].

Bacterial rheotaxis is a passive, physical effect that results from the interplay between self-propulsion, fluid shear, and boundary effects arising from nearby surfaces [4, 9]. Swimming bacteria accumulate near walls [10], where the counter-rotation of the cell body and flagella makes them swim in circles [11]. In shear flows that exist near walls, the body experiences more friction with the wall than the flagella, which sit farther from the surface in faster flow, so the flow swings the flagella downstream and turns the cell upstream, an effect known as weathervaning [2, 4]. The helical shape of the flagella produces an additional torque that turns cells toward the vorticity direction, causing them to drift across the flow [9, 12]. Because these torques grow with shear rate while circling does not, near-surface swimming changes with shear rate to go from circling to upstream swimming, and then to oscillatory motion [4].

Much of our current understanding of rheotaxis comes from changing the flow, through its shear rate and geometry, or the fluid, through its rheology [3, 4, 13–15]. Although faster and straighter swimmers are known to move upstream more effectively [3, 7], much less is known about how the swimmer’s own propulsion apparatus shapes its response to flow. Peritrichous bacteria such as *Escherichia coli* swim using several flagella distributed randomly over the cell body [16–19].

Flagellar number varies between cells, scales with cell length [20], and is tightly regulated [17, 21, 22]. Yet additional flagella increase swimming speed only modestly [23], and speed appears to saturate above approximately five flagella [21]. The drag of the flagellar bundle relative to that of the cell body enters both the weathervane and the chiral torques [4, 9], so flagellar number could potentially affect rheotaxis in ways not captured simply by its effect on swimming speed. However, whether the number of flagella a cell has influences its rheotaxis remains unknown.

Here, we combine experiments and physical modeling to determine how flagellar number shapes bacterial rheotaxis near surfaces. Using an *E. coli* strain with inducible flagellar expression [21], we varied the mean flagellar number from approximately one to six per cell and tracked near-surface swimming across a range of wall shear rates. Increased flagellation reduced downstream transport, increased the fraction of tracks moving upstream, prolonged upstream episodes, and enhanced directed migration across the flow. A model of near-surface swimming, in which flagellar number affects both the swimming speed and the tilt of the cell toward the surface, reproduces the changes in downstream transport, upstream swimming, and lateral migration, linking them to the balance between propulsion and weathervane reorientation. These results identify flagellar number as a property of the cell that shapes its transport in near-surface flows, and suggest a benefit of multiflagellarity that is not revealed by measurements of swimming speed alone.

## Results

### Increased flagellation enhances upstream swimming

To test how the number of flagella affects bacterial motion under flow, we used an *Escherichia coli* strain with IPTG-inducible flagellar expression [21]. Counting flagella by transmission electron microscopy (TEM) showed that the mean flagellar number increased from approximately one per cell without induction to approximately six at 100 *µ*M IPTG (Fig. 1b), although some cells lacked flagella at every IPTG level (Fig. S1). We then tracked cells swimming near the bottom surface of a microfluidic channel, with flow directed in the +*x* direction (Fig. 1a).

**Figure 1:**
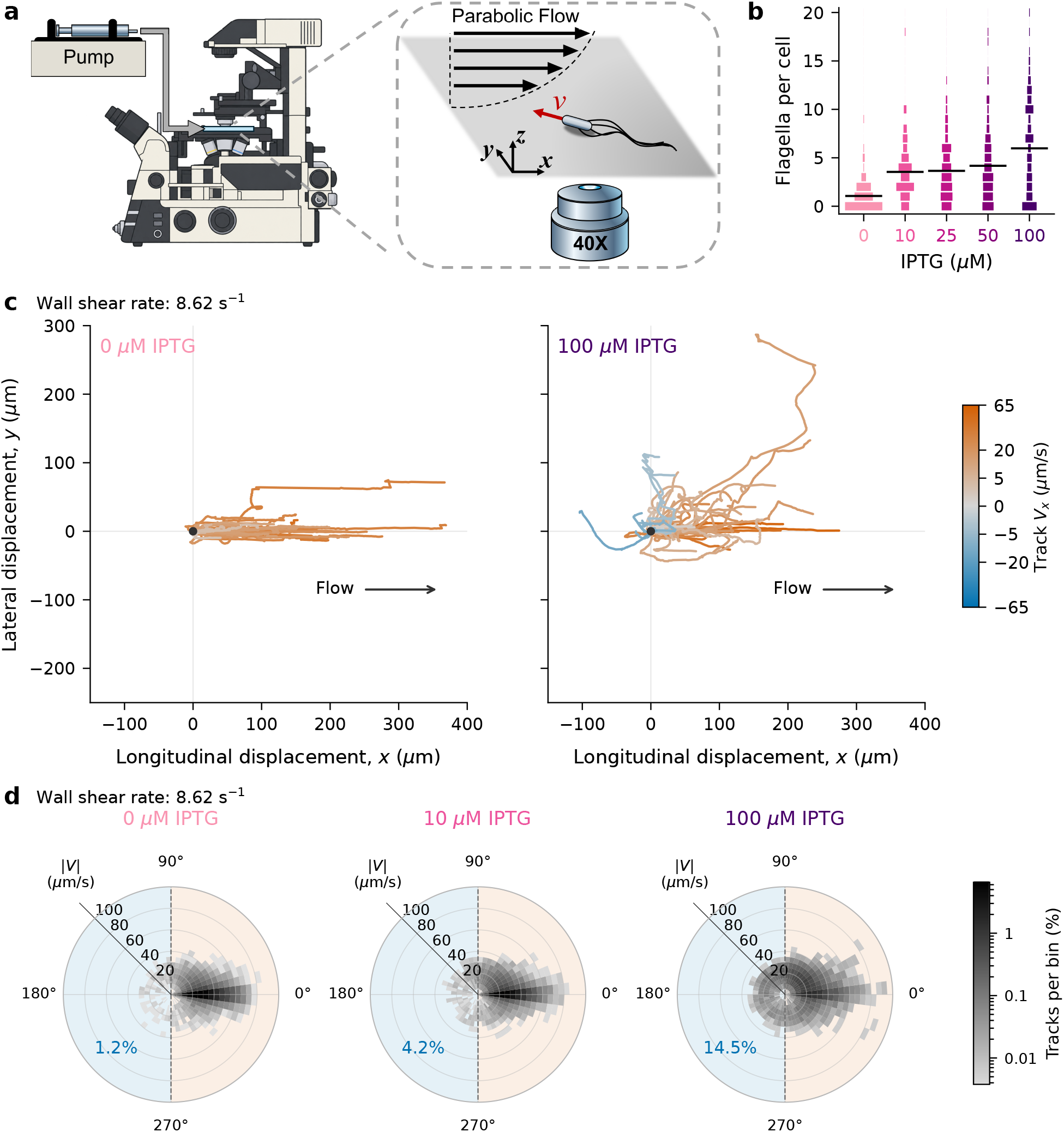
Experimental system and the effect of flagellation under flow. (a) Schematic of the microfluidic setup, imaging plane, flow direction, and coordinates. (b) Distributions of flagella per cell at each IPTG concentration from TEM counts of 120 cells per concentration. Bar length shows the percentage of cells; black lines mark the means. (c) Trajectories of 50 randomly sampled cells per condition at 0 and 100 *µ*M IPTG and a wall shear rate of 8.62 s^−1^. Starting positions are aligned at the origin. Color indicates track-level longitudinal velocity, *V*_*x*_, using a shared signed square-root scale; negative values indicate upstream motion. Arrows indicate the flow direction. (d) Polar heatmaps at 0, 10, and 100 *µ*M IPTG and 8.62 s^−1^. Angle indicates the circular mean of laboratory-frame motion directions within each track, and radius indicates the median speed along that track in *µ*m/s. Color shows the percentage of tracks per bin on a shared logarithmic scale, normalized separately for each condition. Bins span 7.5^°^ and 5 *µ*m/s; blue and orange shading mark the upstream and downstream halves, separated by a dashed line. Numbers give the upstream-track fraction (mean of three biological replicates; Fig. 2b).

Increasing flagellar number enabled more cells to swim against the imposed flow. At the same shear rate, cells with fewer flagella moved predominantly downstream, whereas cells with more flagella displayed more upstream trajectories (Fig. 1c). The joint distributions of swimming speed and direction show this difference across IPTG conditions: increasing IPTG from 0 to 100 *µ*M shifted the distribution toward upstream motion (Fig. 1d). Upstream-directed tracks in the 100 *µ*M IPTG condition reached median speeds as high as 30 *µ*m/s, of the same order as their swimming speeds without flow (Fig. S2). Increased flagellation also enhanced lateral cell migration, resulting in a pronounced bias toward motion in the +*y* direction (Fig. 1c). Thus, changing flagellar number altered both the ability of cells to advance against the current and their migration across it.

For each track, we defined the longitudinal velocity, *V*_*x*_, as the median of its instantaneous velocities along the flow (Methods). Although the population median *V*_*x*_ increased with shear rate at all IPTG concentrations, it was lower in more flagellated populations at any given shear rate (Fig. 2a). This difference became more pronounced at higher shear rates. At the highest shear rate tested (12.31 s^−1^), cells induced with 100 *µ*M IPTG moved downstream approximately 40% more slowly than uninduced cells. Thus, while greater flagellation generally increased swimming speed in the absence of flow (Fig. S2), it reduced downstream transport when flow was imposed.

**Figure 2:**
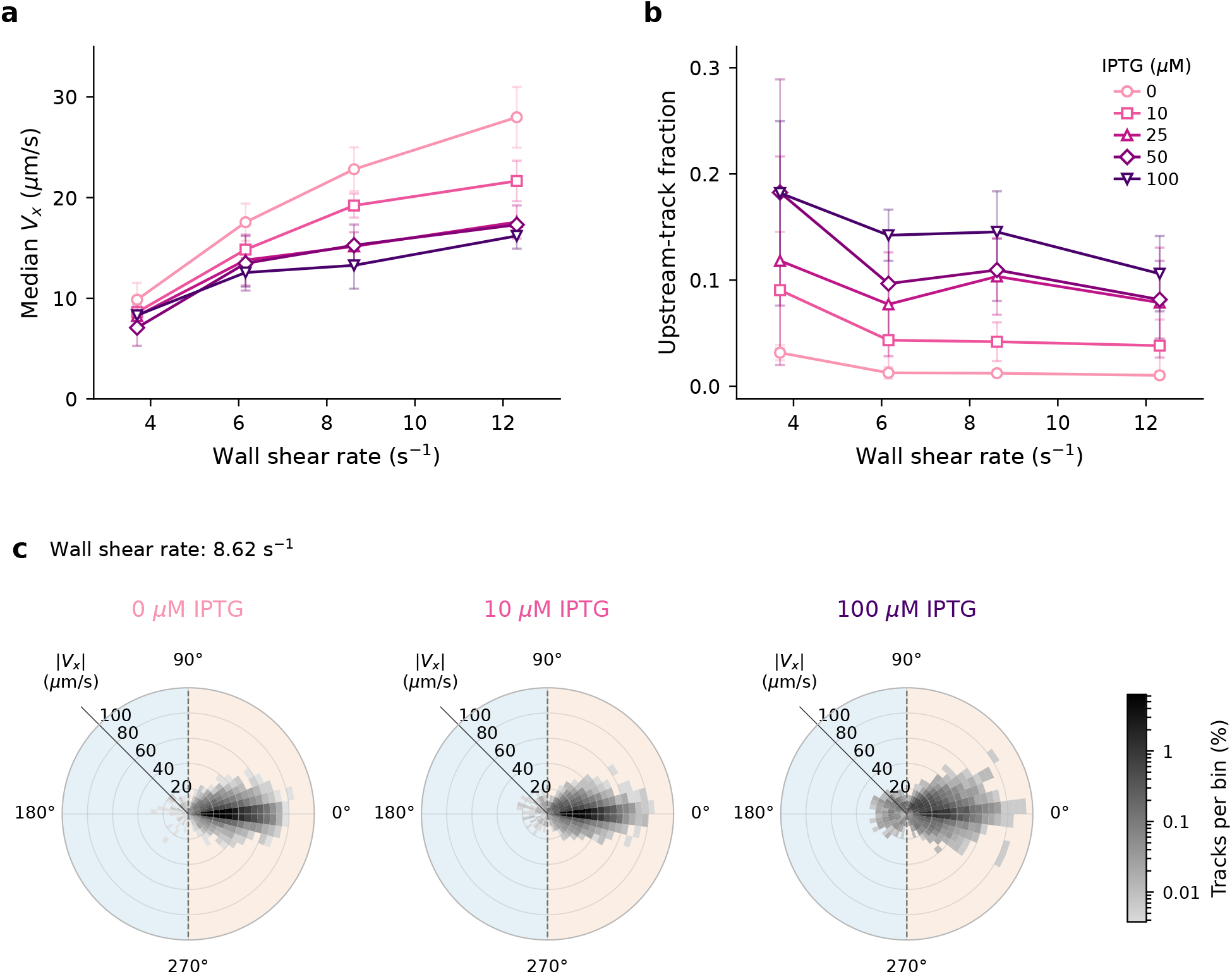
Greater flagellation reduces downstream transport and increases the upstream-track fraction. (a) Population median *V*_*x*_ versus wall shear rate. (b) Upstream-track fraction, the fraction of motile tracks with negative *V*_*x*_, versus wall shear rate. In (a,b), open symbols and error bars show the mean and standard deviation across three biological replicates per condition, for IPTG concentrations of 0, 10, 25, 50, and 100 *µ*M. (c) Polar heatmaps at 0, 10, and 100 *µ*M IPTG and a wall shear rate of 8.62 s^−1^. Angle indicates the circular mean of laboratory-frame motion directions within each track; radius indicates the absolute track-level longitudinal velocity, | *V*_*x*_| . Color shows the percentage of tracks per bin on a shared logarithmic scale, normalized separately for each condition. Bins span 7.5^°^ and 5 *µ*m/s; blue and orange shading mark the upstream and downstream halves, separated by a dashed line.

Increased flagellation also increased the fraction of cells swimming upstream. We classified tracks as upstream when their *V*_*x*_ was negative. In addition to the reports in earlier studies that upstream-track fraction among motile tracks decreased with shear rate [4, 7], we observed that the upstream-track fraction increased with IPTG concentration (Fig. 2b). For instance, at a shear rate of 8.62 s^−1^, increasing IPTG from 0 to 100 *µ*M increased the upstream-track fraction from 1.2% to 14.5%, about a twelvefold increase. The polar distribution of *V*_*x*_ extended farther upstream at higher IPTG concentrations, while remaining approximately symmetric about the flow axis (Fig. 2c). The upstream-track fraction generally continued to increase above 25 *µ*M IPTG, even though the population median longitudinal velocity had largely leveled off in this range (Fig. 2a,b). These observations demonstrate that more flagellated populations resist downstream transport more effectively and contain a larger subpopulation of upstream swimmers, even though their population median velocity remained downstream.

### Increased flagellation enhances directed lateral migration

Greater flagellation enhanced directed migration across the flow. Lateral motion became increasingly asymmetric as flagellar number rose, favoring the +*y* over the *−y* direction across most shear rates, despite the absence of an imposed lateral flow. A one-sided drift near surfaces has been reported before [2, 4, 7], but not its dependence on flagellar number. Representative lateral-displacement traces at 8.62 s^−1^ illustrate this difference between uninduced cells and cells induced with 100 *µ*M IPTG (Fig. 3a). Some trajectories initially moved toward *−y* before reversing their lateral motion toward +*y*. The population median of *V*_*y*_, the median lateral velocity of each track, was close to zero at the lowest shear rate but increased with shear rate, with a stronger increase in more flagellated populations (Fig. 3b). At the highest shear rate tested (12.31 s^−1^), increasing IPTG from 0 to 100 *µ*M increased the population median *V*_*y*_ from approximately 0.2 to 4.7 *µ*m/s (Fig. 3b). The dependence on IPTG concentration largely leveled off above 25–50 *µ*M.

**Figure 3:**
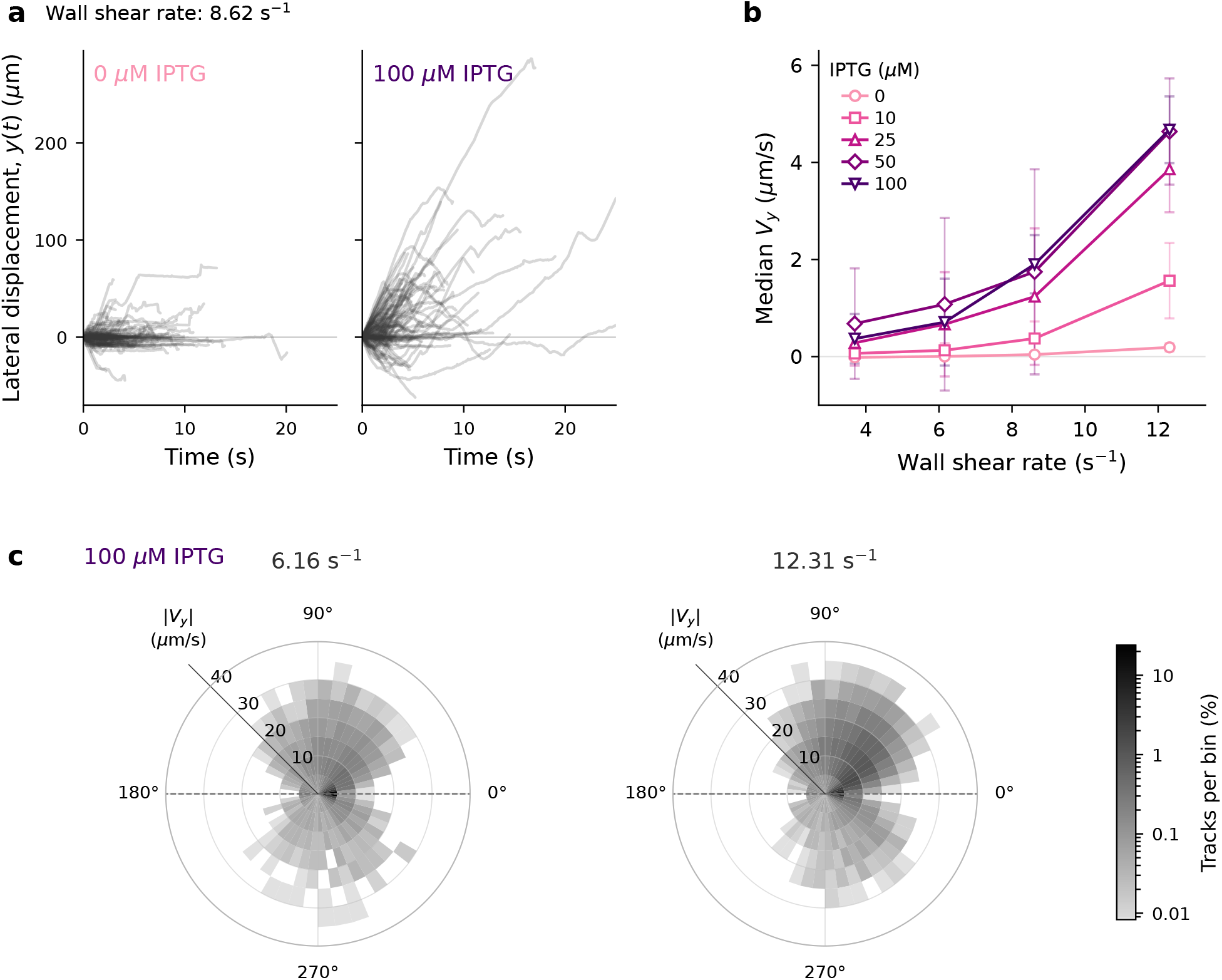
Greater flagellation enhances directed lateral migration. (a) Lateral displacement, *y*(*t*), for 120 randomly sampled tracks per condition at 0 and 100 *µ*M IPTG and a wall shear rate of 8.62 s^−1^, from one biological replicate. (b) Population median lateral velocity, *V*_*y*_, versus wall shear rate for each IPTG concentration. Open symbols and error bars show the mean and standard deviation across three biological replicates per condition. (c) Polar heatmaps at 100 *µ*M IPTG and wall shear rates of 6.16 and 12.31 s^−1^. Angle indicates the circular mean of laboratory-frame motion directions within each track; radius indicates the absolute track-level lateral velocity, |*V*_*y*_|. Color shows the percentage of tracks per bin on a shared logarithmic scale, normalized separately for each condition. Bins span 7.5^°^ and 5 *µ*m/s; the dashed line marks the flow axis and separates the +*y* and *−y* halves.

Unlike the approximately symmetric polar distributions of longitudinal velocity, the lateral-velocity distributions were strongly asymmetric about the flow axis. Stronger flow also increased the asymmetry of lateral motion towards the +*y* direction. At 100 *µ*M IPTG, the polar distributions showed a more pronounced asymmetry at 12.31 than at 6.16 s^−1^, indicating that stronger flow reinforced the preference for +*y* motion while also increasing the magnitude of lateral velocities (Fig. 3c). Thus, while the upstream-track fraction declined with shear rate, directed lateral motion strengthened.

### Increased flagellation prolongs upstream episodes and shortens downstream episodes

To determine how flagellation affects the persistence of motion along the flow axis, we divided trajectories into upstream and downstream episodes according to the sign of their longitudinal velocity and measured each episode’s duration (Fig. 4a). Increasing flagellation shifted upstream episode durations toward longer times and downstream durations toward shorter times (Fig. 4b). This pattern generally held across the tested shear rates (Fig. 4c; additional distributions in Fig. S3a). Nevertheless, downstream episodes remained longer on average than upstream episodes across the measured conditions. Thus, while increased flagellation did not make upstream swimming the longer-lived state, it shifted the balance toward longer upstream episodes.

**Figure 4:**
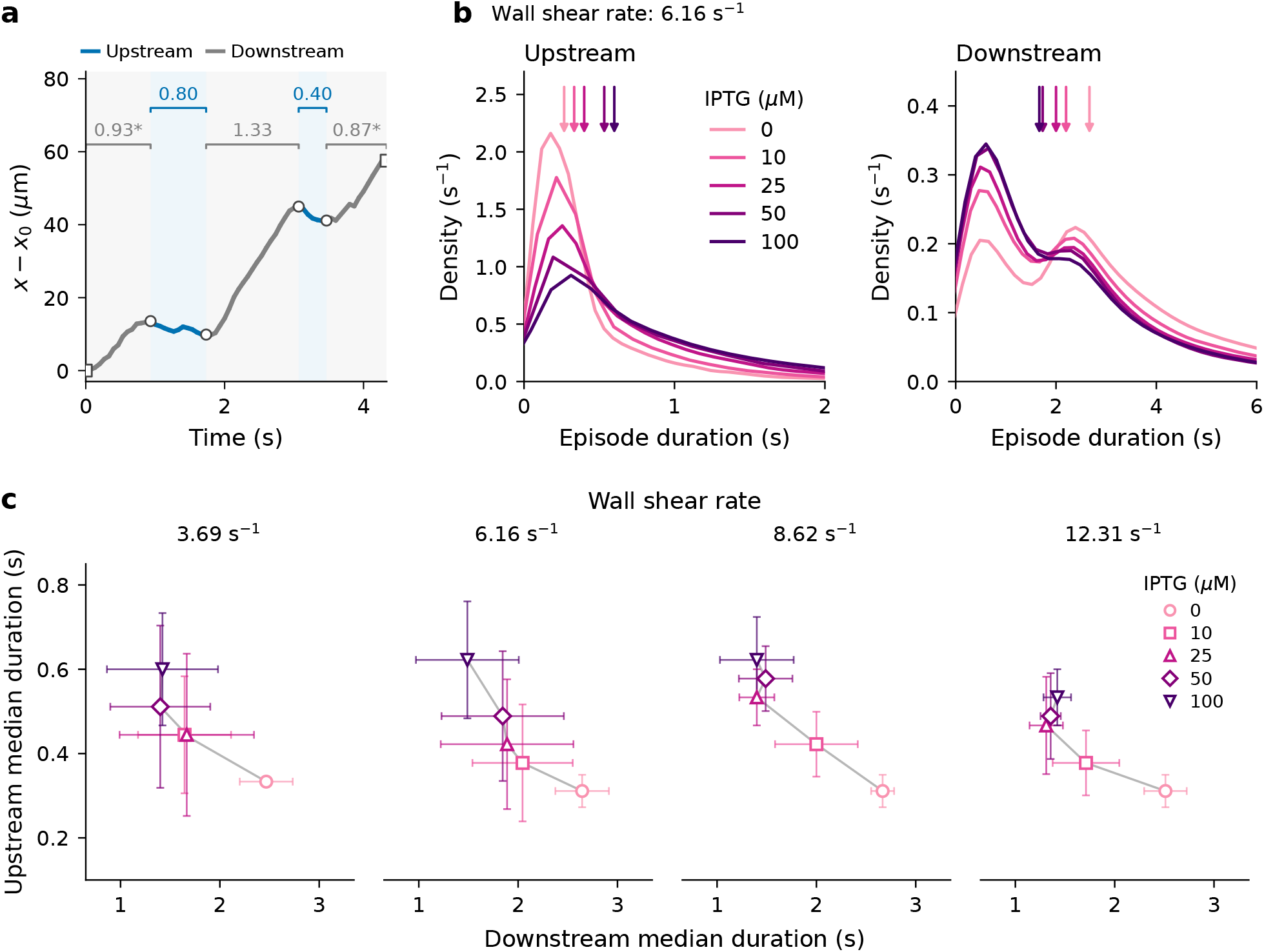
Flagellation alters upstream and downstream persistence. (a) Longitudinal displacement of an example cell at 50 *µ*M IPTG and a wall shear rate of 6.16 s^−1^, colored by upstream (blue) and downstream (gray) classification. Open circles mark direction switches; brackets show episode durations in seconds. Asterisks denote observed durations of episodes intersecting the start or end of the track. These boundary episodes are included in the analysis. (b) Kernel density estimates of upstream and downstream episode durations at a wall shear rate of 6.16 s^−1^ for 0, 10, 25, 50, and 100 *µ*M IPTG. Downward arrows mark the medians of the pooled episode durations. (c) Upstream versus downstream median episode duration at four wall shear rates. Symbols show the means of replicate medians, and horizontal and vertical error bars show the corresponding standard deviations across three biological replicates. Additional duration distributions and their dependence on shear rate are shown in Fig. S3.

Persistence changed little with shear rate over the range tested, and the effect of flagellation held at every shear rate (Fig. 4c; Fig. S3b). At 100 *µ*M IPTG, for example, the mean of the replicate median upstream durations was 0.60–0.62 s from 3.69 to 8.62 s^−1^ and 0.53 s at 12.31 s^−1^, compared with 0.31–0.33 s for uninduced cells. Episodes at the beginning or end of a track are not bounded by direction switches, so we repeated the analysis using only episodes bounded by switches at both ends (Fig. S4). Excluding incomplete episodes removed the second peak in the downstream-duration distribution and substantially reduced downstream median durations, but the upstream durations remained largely unchanged. With complete episodes only, downstream durations differed little between IPTG levels. Nevertheless, increased flagellation remained associated with longer upstream episodes, and downstream episodes remained longer overall than upstream episodes. Thus, including incomplete episodes affected the downstream comparison but preserved the increase in upstream persistence associated with greater flagellation.

### A physical model links flagellation to upstream and lateral motion

To examine how flagellation could influence both upstream swimming and lateral migration, we constructed a stochastic model of bacterial motion near a surface. Cells were advected by the imposed flow (Methods) while swimming at speeds measured in the absence of flow. Their swimming directions evolved through weathervane reorientation [2, 7, 13], near-surface circular swimming [11], chirality-induced turning [4, 9, 12], rotational diffusion [4, 24], and tumbling [24, 25]. Weathervaning turns a cell that is tilted toward the surface to face upstream, because its flagella sit in faster flow than its body; both weathervaning and circling weakened as the gap between the cell body and the surface increased. Each cell kept a fixed height above the surface and a fixed tilt. Flagellar number entered the model in two ways: through the measured swimming speed of each population, and through a prescribed tilt that increased linearly with flagellar number, which strengthened weathervaning. We fitted the model to the median longitudinal and lateral velocities and the upstream-track fraction at 0 and 100 *µ*M IPTG, and then simulated 10, 25, and 50 *µ*M IPTG without refitting (Methods).

The model reproduced the trends in the data (Fig. 5b–e). At a shear rate of 8.62 s^−1^, simulated cells with few flagella were mostly carried downstream, whereas cells with more flagella more often swam upstream and moved farther across the flow (Fig. 5b), as in the experiments (Fig. 1c). Across shear rates, as flagellar number increased, simulated populations were carried downstream more slowly, a larger fraction of tracks moved upstream, and lateral velocity toward +*y* increased (Fig. 5c–e). Stronger flow reduced the upstream-track fraction and increased lateral velocity, as in the experiments. The model predicted these trends at 10, 25, and 50 *µ*M IPTG without refitting (Fig. S5).

**Figure 5:**
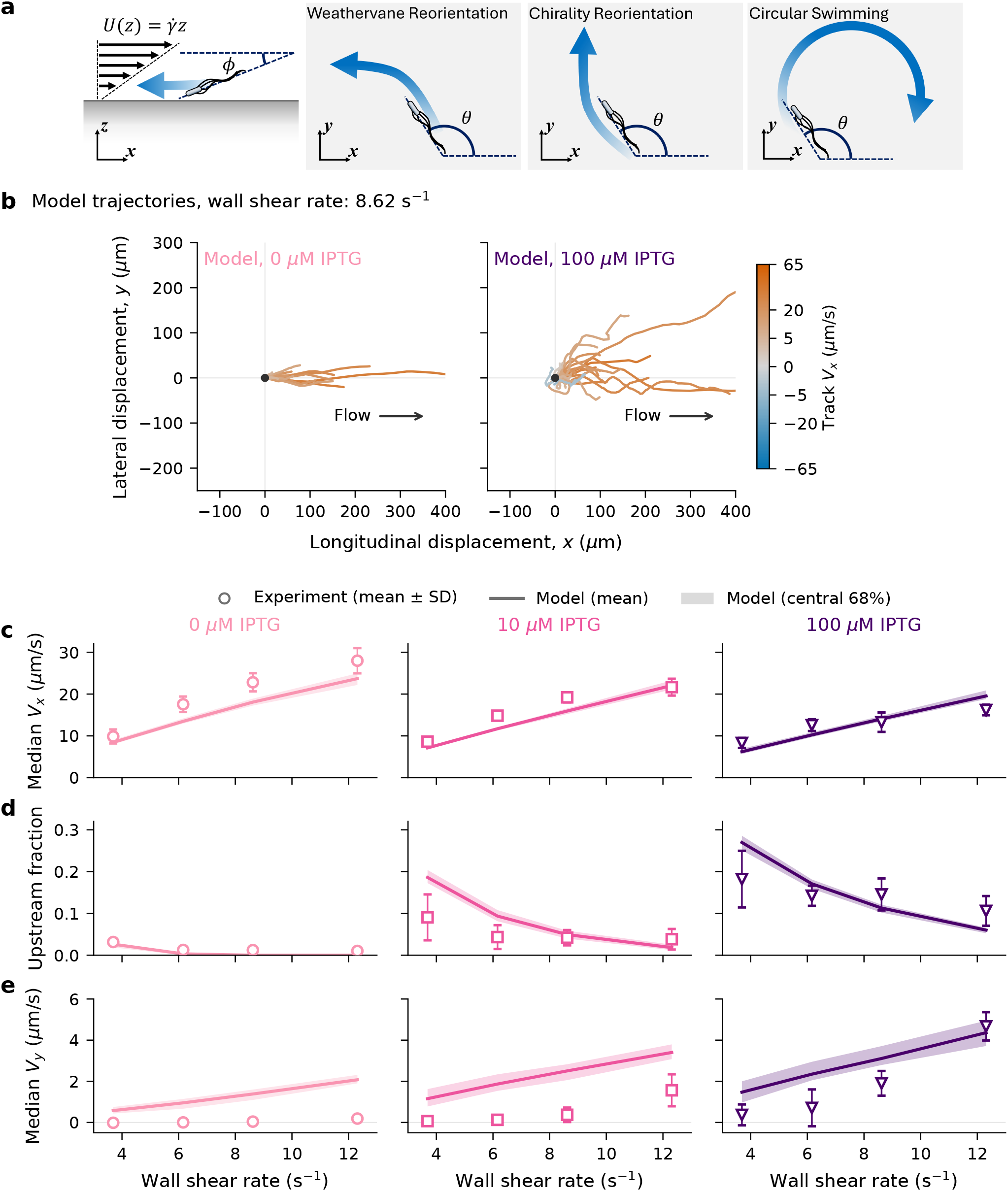
A physical model links flagellation to longitudinal and lateral transport. (a) Model schematic. Side view: a cell tilted toward the surface by *ϕ* in near-wall shear flow. Top views: weathervane reorientation, chirality-induced reorientation, and circular swimming change the heading *θ*. (b) Simulated trajectories at 0 and 100 *µ*M IPTG and a wall shear rate of 8.62 s^−1^. Tracks start at the origin, are cut to durations drawn from the measured track durations, and are colored by track-level longitudinal velocity *V*_*x*_ on the scale of Fig. 1c. (c–e) Median longitudinal velocity *V*_*x*_ (c), upstream-track fraction (d), and median lateral velocity *V*_*y*_ (e) versus wall shear rate at 0, 10, and 100 *µ*M IPTG. Symbols: experiment, mean *±* SD of three biological replicates. Lines and bands: mean and central 68% of 20 independent simulated populations of 200 cells. Parameters were fitted at 0 and 100 *µ*M IPTG.

## Discussion

We varied flagellar number in *E. coli* from approximately one to six flagella per cell and found that greater flagellation changed near-surface rheotaxis in four ways. More flagellated populations were transported downstream more slowly, contained a larger fraction of upstream-swimming cells, maintained upstream motion for longer, and migrated more strongly across the flow. A physical model in which flagellar number affects both the swimming speed and the tilt of cells toward the surface reproduced the trends in downstream transport, upstream swimming, and lateral migration. Flagellation changed the fraction of cells moving upstream more than it changed the median downstream velocity. Above 25 *µ*M IPTG, the population median longitudinal velocity changed little, yet the upstream-track fraction continued to rise. A lower downstream velocity of a population could reflect stronger propulsion, more upstream orientation, or swimming closer to the wall, where the flow is slower [26]. A change in the upstream-track fraction without a corresponding change in the median velocity may reflect a change in how cells orient relative to the flow, without an increase in the upstream velocity of the whole population. These explanations are not exclusive. A faster cell with the same orientation is also more likely to move upstream. In our model, both routes contribute: flagellar number determines the swimming speed and the tilt, and the tilt strengthens weathervane reorientation.

Upstream episodes were longer in more flagellated populations at every shear rate tested, although downstream episodes remained longer overall. The upstream-track fraction and the duration of upstream episodes act together. Cells that reorient upstream more often and stay upstream longer travel farther against the flow before being carried back, much like the cells with long upstream runs that lead contamination of channels [8]. The combined effect on how far cells travel upstream may therefore exceed what either measurement suggests alone.

Flagellation also strengthened directed migration across the flow. The bias toward +*y* is consistent with the sideways drift reported for *E. coli* near surfaces [2, 7] and identified as drift toward the vorticity direction by Mathijssen et al. [4]. In our geometry, +*y* was the positive vorticity direction near the bottom surface, where we recorded images; on the top surface, where vorticity changes sign, the drift would be toward *−y*. The increase in lateral drift with flagellar number could reflect a change in the chiral torque [9, 12]. In our model, however, the chiral strength was kept constant, yet the simulated lateral velocity still increased with flagellar number (Fig. 5e). Changes in chiral torque associated with flagellation are therefore not required to explain the enhanced lateral migration.

These results suggest one reason cells might invest in flagella beyond the number needed for maximal speed. Wild-type *E. coli* grown in rich medium sits near the speed plateau [21], yet flagella added in this range would still improve the ability of cells to hold position and advance against flow. Variation in flagellar number between cells [20, 21] would also produce variation in rheotactic ability within a clonal population, so that the most flagellated cells could lead upstream spread, much as non-genetic diversity shapes chemotactic migration [27]. The shear rates in our experiments, 3.7 to 12.3 s^−1^, are comparable to those in urinary catheters under steady drainage (about 3 s^−1^) [4], where upstream swimming may contribute to catheter-associated infection.

Several extensions of this work would sharpen these conclusions. We tracked cells in two dimensions without resolving their height above the surface. We also imaged near a flat wall away from channel edges. Channel edges can strengthen upstream swimming [2, 3, 14], and whether flagellar number amplifies or saturates these effects remains to be tested. Measuring orientation and height directly [4, 25, 28], together with flagellar number on individual cells, would separate the contributions of propulsion and alignment that our population measurements can only infer.

## Methods

### Strain and growth conditions

We used *Escherichia coli* strain VS1683 (*Ptac*), in which the chromosomal flagellar master regulator operon *flhDC* is controlled by the IPTG-inducible *P*_tac_ promoter [21]. A copy of *lacI* upstream of the promoter and an additional copy supplied by the pTrc99a plasmid reduce basal expression. IPTG relieves repression by LacI, allowing flagellar expression to be varied. Cells were grown overnight at 37 ^°^C in LB with 100 *µ*g/mL ampicillin. Overnight cultures were diluted 1:100 into tryptone broth (TB) containing 100 *µ*g/mL ampicillin and 0, 10, 25, 50, or 100 *µ*M IPTG, and grown at 37 ^°^C with shaking at 200 rpm to an OD_600_ of 0.45–0.55.

### Flagellar counting

Flagella were counted on cells imaged by transmission electron microscopy (TEM). Cultures were grown to an OD_600_ of 0.45–0.55 as described above. Copper TEM grids (400 mesh, carbon film; Electron Microscopy Sciences #CF400-Cu-50) were glow-discharged for 60 s. One milliliter of culture was centrifuged at 1500 *×g* for 7 min at room temperature, and the pellet was gently resuspended in 980 *µ*L of phosphate-buffered saline and chilled on ice for 5 min. Cells were fixed by adding 20 *µ*L of 50% glutaraldehyde (final concentration 1%) and incubating on ice for 5 min. Ten microliters of cell suspension were placed on a grid for 5 min, and the excess was wicked away with filter paper. Grids were washed twice with ddH_2_O (10 s each), stained twice with 10 *µ*L of UranyLess EM Stain (Electron Microscopy Sciences #22409) for 4 min each, washed twice more (5 s each), and dried at room temperature.

Grids were imaged on a Talos L120C TEM at 120 kV with a Ceta-S camera (4096 *×* 4096 pixels, 14 *µ*m pixel size). For each IPTG concentration, three biological replicate grids were prepared and 40 cells were imaged per grid. Counts from the three grids were pooled (120 cells per concentration) to construct flagellar-number distributions and calculate the mean number of flagella per cell (Fig. 1b; Fig. S1).

### Microfluidic device fabrication

The microfluidic device consisted of a straight channel 5 mm long, 900 *µ*m wide, and 95 *µ*m high, preceded by a serpentine hydraulic resistor 34 mm long and 140 *µ*m wide to stabilize flow. We fabricated the master mold by UV photolithography using SU-8 2035 photoresist (Kayaku Advanced Materials Inc.) on a 10 cm-diameter silicon wafer. Polydimethylsiloxane (PDMS) and curing agent (SYLGARD 184, Dow) were mixed at a 10:1 ratio (w/w), poured on the master mold in a petri dish, degassed under vacuum, and cured overnight at 65 ^°^C. Cast PDMS was peeled off the master mold and cut, and the inlet and outlet ports were punched (Harris Uni-Core 1.0 mm). The PDMS slab was bonded to a 40 *×* 24 mm No. 1 coverslip (Thermo Scientific) after treatment in an oxygen plasma cleaner (PDC-001, Harrick Plasma Inc.) at an RF power of 18 W for 1.5 min. To reinforce bonding, devices were baked at 65 ^°^C for an additional 2 h.

### Flow and imaging

Before introducing cells, we flushed the channel with motility buffer (6.15 mM K_2_HPO_4_, 3.85 mM KH_2_PO_4_, 0.1 mM EDTA, 66.6 mM NaCl, 1% glucose, 0.001% Tween-80, pH 7.0) to remove air bubbles, limit cell attachment to the glass, and support near-surface swimming. For each experiment, 1 mL of culture was centrifuged at 1500 *×g* for 7 min, and the pellet was resuspended in 1 mL of motility buffer. Cells were loaded into a 1 mL plastic syringe (BD) mounted on a syringe pump (NE-500, New Era Pump Systems) and introduced through PE/2 tubing (Scientific Commodities Inc.). Flow rates were 0.3, 0.5, 0.7, and 1.0 *µ*L/min, corresponding to wall shear rates of 3.69, 6.16, 8.62, and 12.31 s^−1^, respectively. Wall shear rates at the imaging region (*z* = 0, *y* = *W/*2) were calculated as 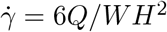, assuming a flow profile that is parabolic in *z* and flat in *y, u*_*x*_(*z*) = 6*Qz*(*H − z*)*/WH*^3^.

Cells were imaged near the bottom surface of the channel in the *xy* plane. The +*x* direction followed the imposed flow, *y* denoted the lateral direction, and *z* was perpendicular to the channel surface. The 250 *×* 250 *µ*m region of interest was centered across the channel width (Fig. 1a), at about 3 mm downstream from the main channel opening. Phase-contrast recordings were acquired at 15 frames/s using an ORCA-Fusion BT CMOS camera on a Nikon Ti2E microscope equipped with a Nikon CFI Plan Fluor DLL 40*×*, NA 0.75 objective. Each recording lasted 45 s, and 4 recordings were made for each of the 3 biological replicates of each condition. Measurements were performed at room temperature, after allowing about 1 min following each change in flow rate.

Swimming speed without flow was measured near the glass surface in the same microfluidic channels, using cells from the same cultures and the same growth, IPTG, and imaging conditions as the flow experiments. Measurements included three biological replicates for each of the five IPTG concentrations, four recordings per replicate, and more than 3000 tracks per replicate (Fig. S2). The mean of the replicate median speeds at each IPTG concentration (Fig. S2) was 5.4, 11.3, 14.6, 18.6, and 17.5 *µ*m/s; the model used 5, 12, 15, 18, and 17 *µ*m/s.

### Trajectory analysis

Recorded ND2 files were processed in Fiji [29] using TrackMate [30] to identify and link cells across frames. A Jython script with a template TrackMate settings file was used to automate the process of spot detection and track assignment. Each file was pre-processed with a 3 *×* 3 median filter, and a difference of Gaussian (DoG) detector was implemented for a sub-pixel localization of spots with an estimated object diameter of 3.6 *µ*m. To optimize the selected quality threshold for each file, the histogram of quality threshold values aggregated from the first 3 frames was processed with Otsu’s method to leave out spurious detections. To connect the detected spots, TrackMate simple LAP tracker was used with a maximum linking distance of 10 *µ*m, and a maximum gap-closing distance of 10 *µ*m. A maximum frame gap of 3 was used to link missing spots across frames.

Tracks shorter than 2 s were discarded. Tracks with insufficient motion were also excluded if the variance of sum of their coordinates Var(*X* + *Y*) calculated over their track frames was smaller than 4 *µ*m^2^.

Instantaneous longitudinal and lateral velocities were calculated from cell positions using the central-difference method. Track-level swimming speed was defined as the median of instantaneous speed. Longitudinal and lateral velocities were summarized within each track using median of their instantaneous velocities in each direction respectively. Tracks with negative median longitudinal velocity were classified as upstream, and those with positive median longitudinal velocity as downstream. The upstream-track fraction was calculated as the number of upstream tracks divided by the total number of motile tracks (defined below).

The mean direction of each track was calculated as the circular mean of its instantaneous motion directions, 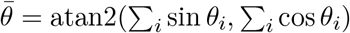, where *θ*_*i*_ is the velocity direction at frame *i*. Each direction contributed equally, irrespective of speed. Circular variance was calculated as 1 *− R*, in which the mean resultant length 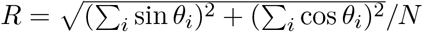 [31]. To exclude the washed-out cells with least motility relative to flow direction, a nonmotile track was defined as a track with absolute value of circular mean smaller than 0.05 rad while having a circular variance smaller than 0.002. This definition was not implemented in the no-flow conditions. Polar heatmaps shown in Figs. 1d, 2c, and 3c combined the circular mean direction with the track-level speed or absolute longitudinal or lateral velocity. Bins spanned 7.5^°^ and 5 *µ*m/s. Color represented the percentage of tracks in each bin, normalized separately for each condition and displayed on a shared logarithmic scale within each comparison.

Population velocity summaries were calculated as the median of track-level measurements within each biological replicate. Reported values and error bars show the mean and standard deviation across three biological replicates. Upstream-track fractions were calculated separately for each replicate before averaging. Representative trajectory plots used randomly sampled tracks from one biological replicate, with sample sizes specified in the figure captions.

### Persistence analysis

Trajectories were divided into upstream and downstream episodes according to the sign of longitudinal motion. To reduce fluctuations caused by cell-body wobble and localization noise, we applied a centered five-frame moving average to the instantaneous longitudinal velocity, *v*_*x*_. At the first and last two frames of each track, the average used the three or four available frames. Direction switches were identified from changes in the sign of *v*_*x*_, with the additional criterion that *v*_*x*_ change by more than 1 *µ*m/s; otherwise, the previous direction was retained. For tracks that never crossed this threshold, the sign of the smoothed *v*_*x*_ was used. Tracks starting with undecided frames were back-filled from the first later confirmed direction.

Episode durations were calculated using the number of consecutive frames with the same signed direction, including the switching frame, while excluding the missing frames.

The main analysis included incomplete episodes intersecting the beginning or end of a track. These boundaries can arise when cells enter or leave the region of interest, move into or out of focus [25], or remain in view when recording begins or ends. Their observed durations therefore need not represent complete intervals between direction switches. We repeated the analysis after excluding these boundary episodes, retaining only episodes bounded by observed switches at both ends (Fig. S4). Requiring both switches to occur within a track can preferentially exclude long episodes [32], so this comparison does not correct for unobserved portions of episodes.

Upstream and downstream durations were summarized separately. Median durations were calculated within each biological replicate, and summary plots show the mean and standard deviation across three biological replicates. Duration distributions were displayed using kernel density estimates with a Gaussian kernel and bandwidth *h* = *sn*^−1*/*5^, where *s* is the sample standard deviation and *n* is the number of episodes.

### Model

We simulated cells swimming near the bottom surface in the flow 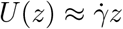, which approximates the parabolic channel flow within the 5 *µ*m imaged layer. Each cell swam at a fixed speed *v*, at a fixed height *z*_*b*_, and with a fixed tilt *ϕ* toward the surface; only its heading *θ* in the *xy* plane changed. Positions evolved as

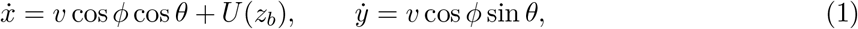

and the heading as

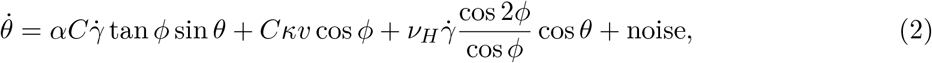

where the three terms describe weathervaning, circular swimming near the surface, and chirality-induced turning, and the noise term includes rotational diffusion and tumbles. The wall-coupling factor

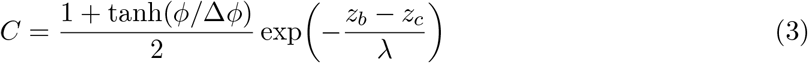

weakens weathervaning and circling as the gap *z*_*b*_ *−z*_*c*_ between the cell body and the surface grows, where *z*_*c*_ is the height of the body center when the body touches the surface.

Flagellar number *N* entered the model in two ways. Each population swam at its measured mean speed (Flow and imaging), and the tilt increased linearly with flagellar number, *ϕ*(*N*) = *ϕ*_0_ + *ϕ*_1_(*N −* 2), with *N* set to the mean number of flagella on flagellated cells (2, 4, 4, 5, and 7 at 0, 10, 25, 50, and 100 *µ*M IPTG). Speed, height, and circling curvature varied across cells. We fitted five parameters (*ϕ*_0_, *ϕ*_1_, *α, λ*, and Δ*ϕ*) to the population median *V*_*x*_ and *V*_*y*_ and the upstream-track fraction at 0 and 100 *µ*M IPTG, and then simulated 10, 25, and 50 *µ*M IPTG without refitting. Simulated tracks were analyzed with the same pipeline as the experimental tracks. Each condition was simulated as 20 independent populations of 200 cells. The full model, parameter values, and fitting procedure are given in the Supplementary Information (Model details; Table S1).

## Acknowledgments

We thank Victor Sourjik for sharing the *Ptac* strain (VS1683), Luis Meneses for help with experiments, Doug Shepherd for helpful discussions, Alexandra Ros for providing microfluidic fabrication equipment, and Mukul Sonker for help in fabrication of microfluidic devices. This work was supported by awards from the Arizona Biomedical Research Centre (grant RFGA2023-008-14), the National Institute of General Medical Sciences of the National Institutes of Health (grant R00GM134124), and the U.S. National Science Foundation (DBI-2119963).

## Author contributions

F.J. designed and performed the experiments, analyzed the data, developed the model, and wrote the manuscript. S.Y. performed electron microscopy and counted flagella. N.W. supervised the project, contributed to modeling and data analysis, and wrote the manuscript.

## Supplementary Information

### Model details

#### Flow

The flow velocity at height *z* above the bottom surface was

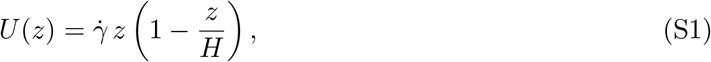

where *H* is the channel height. Because the channel is about ten times wider than it is tall, the flow was taken to be constant across its width.

#### Cell height and speed

Each cell had a constant body height *z*_*b*_ = *z*_*c*_ + *h*, where

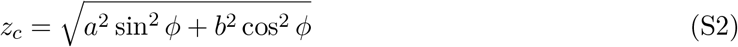

is the height of the body center when the cell touches the surface, *a* = 1 *µ*m and *b* = 0.5 *µ*m are the semi-axes of the prolate spheroid body, and *h* was drawn from an exponential distribution with mean 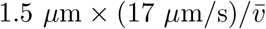, because hydrodynamic attraction to the surface increases with swimming speed [10]. Speeds were drawn from a lognormal distribution with mean 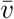 and standard deviation 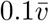.

#### Heading dynamics

For simplicity, the weathervane torque was calculated from the difference in flow velocity between the center of the body and the midpoint of the flagella, and from the distance between them [2, 7]; in a linear shear flow this gives the weathervane term in the Methods. Since the weathervane effect is usually described for a cell body in contact with the surface, the wall-coupling factor *C* weakens it, and the circling term, as the gap between the body and the surface increases or as the cell tilts away from the surface. The curvature *κ* is negative to capture clockwise circling, and its magnitude was drawn for each cell from an exponential distribution with mean |*κ*| = 0.05 *µ*m^−1^.

#### Noise and tumbles

Runs (*χ* = 0) had exponentially distributed durations with mean 1 s. Tumbles (*χ* = 1) stopped swimming for a fixed duration *τ*_*T*_ = 0.1 s. Rotational noise was added to the heading as

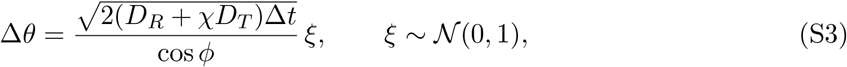

where *D*_*R*_ = 0.057 rad^2^/s is the rotational diffusion coefficient and 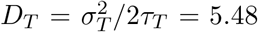 rad^2^/s, with *σ*_*T*_ = 60^°^, is an additional tumble diffusion coefficient applied only during tumbles [4].

#### Fitting

All other parameters were fixed at literature values or set before fitting (Table S1). Simulated cells were cut into tracks with the measured track-length distribution and analyzed with the same pipeline as the experimental tracks. Fits minimized the squared deviations from the measured values, scaled by the replicate standard deviations. Three fitted parameters (*ϕ*_0_, *λ*, and Δ*ϕ*) lay at the bounds of their search ranges.

**Table S1:**
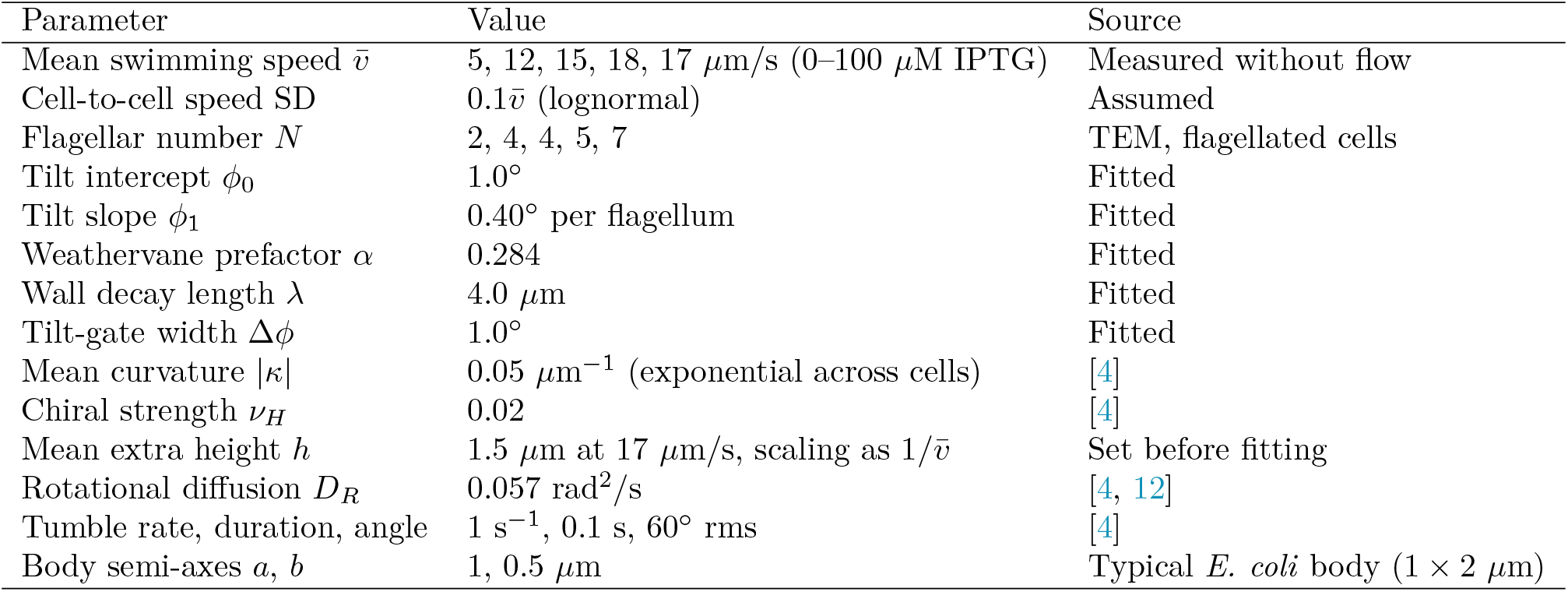
Model parameters.

**Figure S1:**
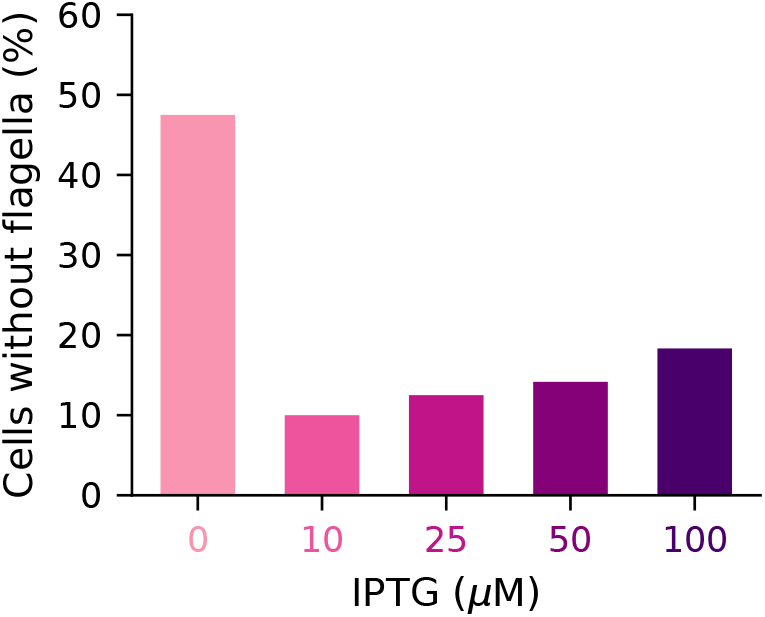
Cells without flagella across IPTG concentrations. Percentage of cells with no flagella at each IPTG concentration, from TEM counts of 120 cells per concentration. The full distributions are shown in Fig. 1b.

**Figure S2:**
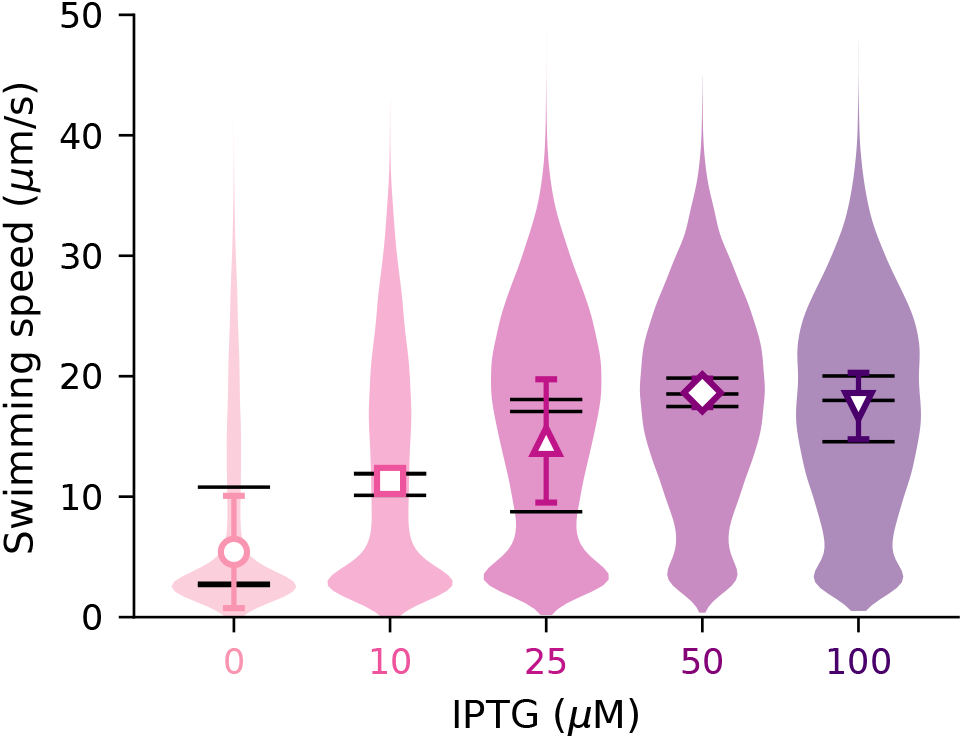
Swimming speed in the absence of flow. Median instantaneous speed of each track near the glass surface without flow. Violins show all tracks at each IPTG concentration, pooled across three biological replicates. Black ticks mark the replicate medians. Symbols and error bars show the mean *±* SD of the replicate medians.

**Figure S3:**
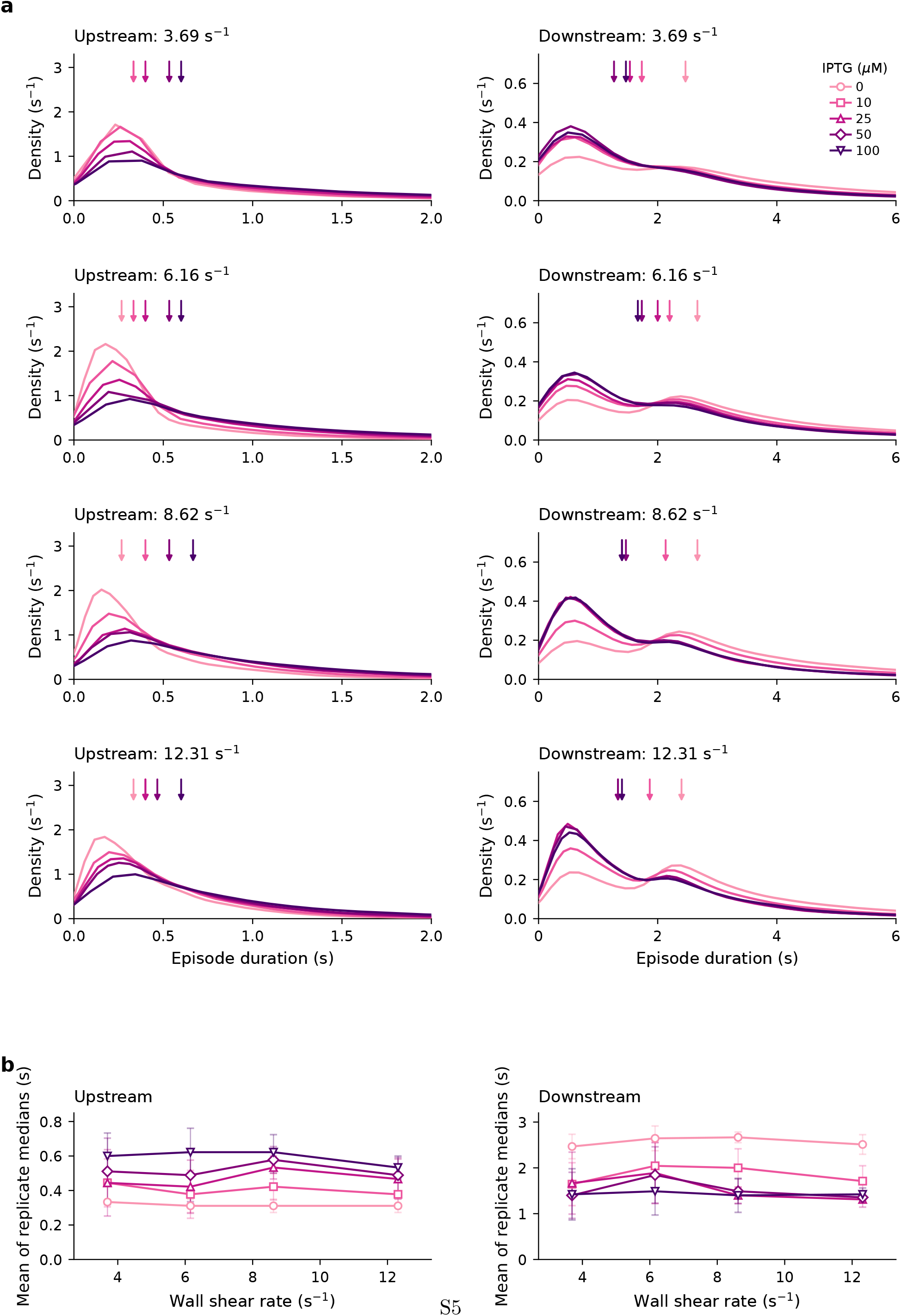
Directional persistence with track-boundary episodes included. Episodes intersecting track starts or ends are included, as in Fig. 4. (a) Kernel density estimates of upstream (left) and downstream (right) episode durations at wall shear rates of 3.69, 6.16, 8.62, and 12.31 s^−1^ for 0, 10, 25, 50, and 100 *µ*M IPTG. Downward arrows mark the medians of the pooled episode durations. Longer tails extend beyond the displayed duration ranges. (b) Upstream and downstream episode durations at all four wall shear rates. Symbols show the means of replicate medians, and error bars show standard deviations across three biological replicates.

**Figure S4:**
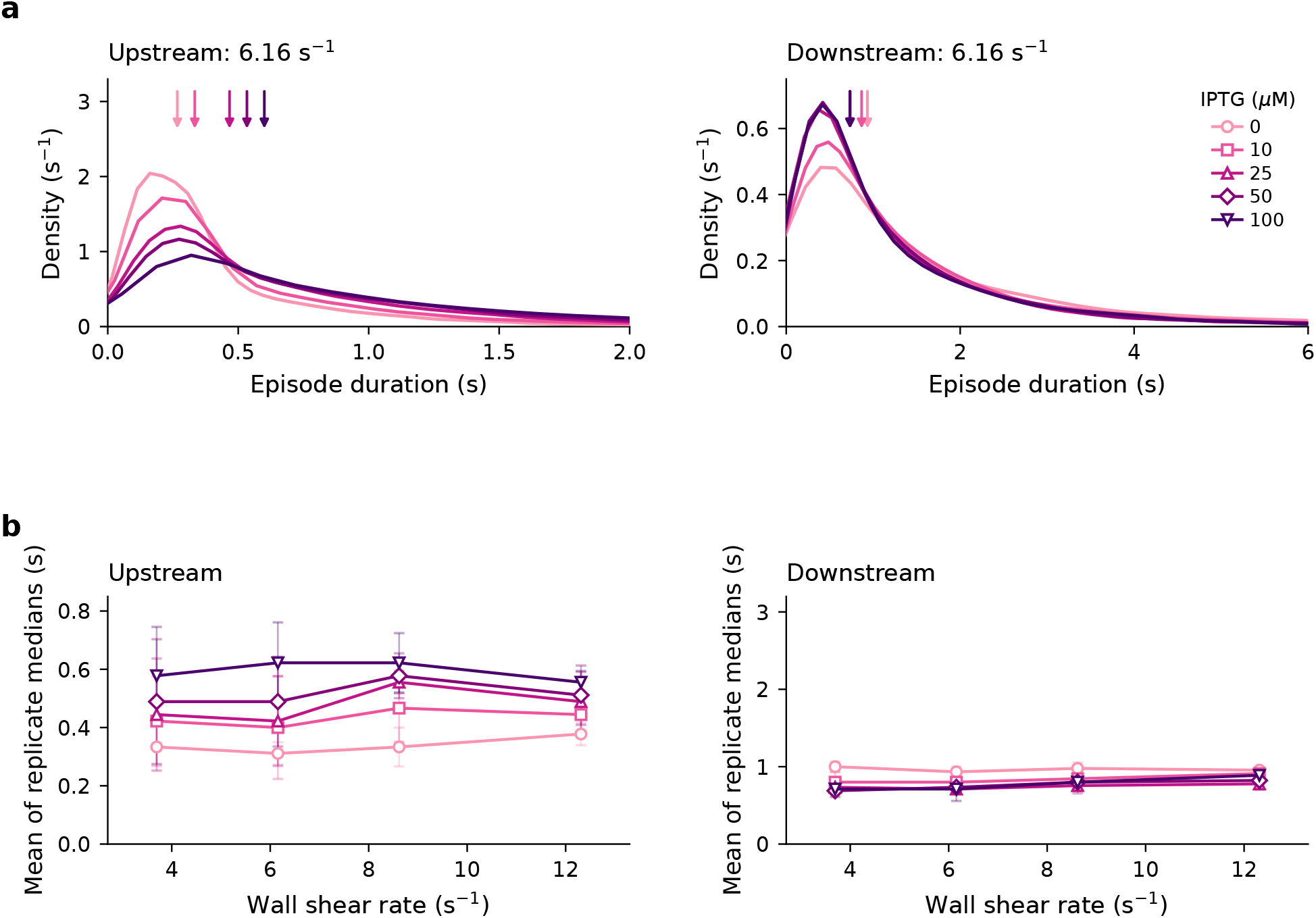
Directional persistence with track-boundary episodes excluded. Only episodes bounded by observed direction switches at both ends are considered. (a) Kernel density estimates of upstream (left) and downstream (right) episode durations at a wall shear rate of 6.16 s^−1^ for 0, 10, 25, 50, and 100 *µ*M IPTG. Downward arrows mark the medians of the pooled episode durations. (b) Upstream and downstream episode durations at all four wall shear rates. Symbols show the means of replicate medians, and error bars show standard deviations across three biological replicates.

**Figure S5:**
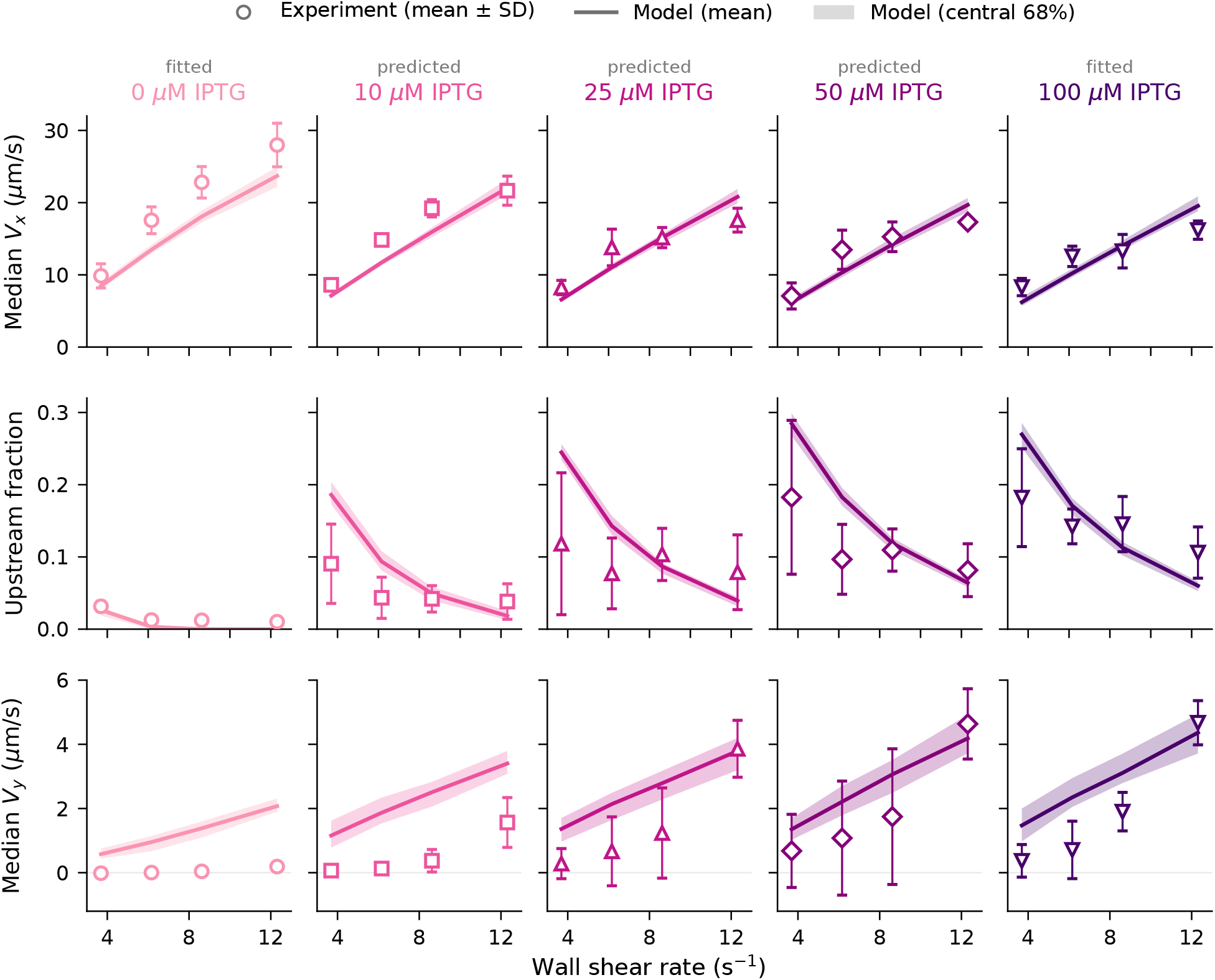
Model and experiment at all IPTG levels. Median longitudinal velocity *V*_*x*_ (top), upstream-track fraction (middle), and median lateral velocity *V*_*y*_ (bottom) versus wall shear rate at 0, 10, 25, 50, and 100 *µ*M IPTG. Symbols: experiment, mean *±* SD of three biological replicates. Lines and bands: mean and central 68% of 20 independent simulated populations of 200 cells. Parameters were fitted at 0 and 100 *µ*M IPTG; 10, 25, and 50 *µ*M are predictions. Simulated tracks at 25 and 50 *µ*M were cut to the measured track durations at 0 *µ*M.

